# Development of the Intestinal Microbiome in Cystic Fibrosis in Early Life

**DOI:** 10.1101/2022.05.27.493808

**Authors:** Courtney E. Price, Thomas H. Hampton, Rebecca A. Valls, Kaitlyn E. Barrack, George A. O’Toole, Juliette C. Madan, Modupe O. Coker

## Abstract

Cystic Fibrosis is a heritable disease that causes altered physiology at mucosal sites; these changes result in chronic infection in the lung, significant gastrointestinal complications as well as dysbiosis of the gut microbiome, although the latter has been less well explored. Here, we describe the longitudinal development of the gut microbiome in a cohort of children with cystic fibrosis (cwCF) from birth through early childhood (0-4 years of age) using 16S rRNA gene amplicon sequencing of stool samples as a surrogate for the gut microbiota. Similar to healthy populations, alpha diversity of the gut microbiome increases significantly with age, but diversity plateaus ∼2 years of age for this CF cohort. Several taxa that have been associated with dysbiosis in CF change with age towards a more healthy-like composition; notable exceptions include *Akkermansia*, which decreases with age, and *Blautia*, which increases with age. We also examined the relative abundance and prevalence of nine taxa associated with CF lung disease, several of which persist across early life, highlighting the possibility of the lung being seeded directly from the gut early in life. Finally, we applied the Crohn’s dysbiosis index to each sample, and found that high Crohn’s-associated dysbiosis early in life (<2 years) was associated with significantly lower *Bacteroides* in samples collected from 2-4 years of age. Together, these data indicate a persisting dysbiosis in the gut microbiota as well as markers associated with inflammatory bowel disease early in life for cwCF.

**IMPORTANCE:** Cystic Fibrosis is a heritable disease that disrupts ion transport at mucosal surfaces, causing a buildup of mucus and dysregulation of microbial communities in both the lungs and the intestines. Persons with CF are known to have dysbiotic gut microbial communities, but the development of these communities over time beginning at birth have not been thoroughly studied. Here, we describe the development of the gut microbiome of cwCF throughout the first four years of life, during the critical window of both gut microbiome and immune development. Our findings indicate a persisting dysbiosis, the possibility of the gut microbiota as a reservoir of airway pathogens and a surprisingly early indication of a microbiota associated with inflammatory bowel disease.

## INTRODUCTION

Cystic fibrosis (CF) is a heritable disease caused by mutations in the cystic fibrosis transmembrane conductance regulator (CFTR). Loss of CFTR function leads to altered secretion of chloride and bicarbonate, and accumulation of abnormally thick mucus in both the lungs and the intestinal tract [1, 2]. Loss of CFTR function also alters bile acid production and diminishes secretion of pancreatic enzymes. Persons with CF (pwCF) can experience intestinal blockages at birth (meconium ileus), bacterial overgrowth in the small bowel (SIBO), inflammation, dysmotility, and often struggle with sufficient weight gain in early childhood [3].

More recently, several studies have documented alterations in the gut microbiome for pwCF, as well as associations between the gut microbiome and health outcomes [4-10]. Alterations in the gut microbiome are associated with, and may contribute to, important clinical outcomes, including increased inflammation, maldigestion, malabsorption, intestinal lesions and poor linear growth [8, 11-14]. Emerging evidence indicates that the gut-lung axis, wherein the health of the intestinal microbiome affects distal organ health, is an important determinant of lung health outcomes for pwCF [5-7], likely via immune programming. Studies have shown that weight gain is associated with better pulmonary outcomes, and that in early life the gut microbiome is a better predictor of lung health outcomes than the respiratory microbiome [15]. Microbiome alterations are driven by CFTR mutations and are therefore inherent to CF [10, 16], but are also likely influenced by CF clinical manifestations such as pancreatic sufficiency and CFTR genotype [17-19]. CF gut microbiome health can also be influenced by external exposures, such as breastfeeding or delivery mode, and by treatment with CF-specific drug regimens, probiotics, or antibiotics [5, 20-23]. However, studies comparing microbiome development in children with and without CF point to the decreased influence of these typical exposures in infancy, highlighting the importance of CFTR mutations and the associated micro- and macro-environments in CF [4, 9].

For pwCF, gut microbiome dysbiosis begins in early childhood and continues into adolescence and adulthood [17, 23]. Infants and children with CF (cwCF) have been shown to have lower alpha diversity [4, 9, 11, 17, 24] as well as delayed microbiome maturation relative to healthy cohorts [8]. Additionally, cwCF have many of the same alterations noted in adults, including increased *Veillonella* and *E. coli*, and lower *Bacteroides, Faecalibacterium*, and *Akkermansia*. However, dysbiosis is most pronounced in early infancy, and some differences between CF and healthy cohorts are age-dependent [8, 17]. Furthermore, an age-based analysis revealed that CF intestinal microbiomes appeared more immature than microbiomes of healthy children, which could have implications for immune development.

The early window of gut microbiome maturation (0-∼4 years) is particularly important because development of the immune system occurs during this same time period, and a healthy gut microbiome is essential for proper immune training [25-27]. This early developmental window has only been fully analyzed in two CF studies, and partially covered by several others [4, 8, 9, 11, 16, 17, 24, 28-30]. CF microbiome studies to date have primarily focused on either the early microbiomes (<1 year), adult microbiomes, and a few small studies have sampled at larger intervals throughout childhood [8, 9, 28]. Here, we characterize fecal microbiomes from a cohort of 39 children across the first 4 years of life, with samples taken frequently throughout this important developmental period.

## RESULTS AND DISCUSSION

### Age Influences the Cystic Fibrosis Microbiome Alpha- and Beta-Diversity

In healthy cohorts, age is the dominant factor shaping the structure of the microbiome [31]. In this study, we wanted to determine whether the intestinal microbiome of cwCF exhibits known maturation patterns, including increases in microbial diversity over time and age-dependent changes in microbial composition. We used a mixed linear model with patient as the random effect to test whether Shannon Diversity Index (SDI), a measure encompassing both microbial richness and evenness, was significantly correlated with age in days for cwCF; when including all samples, we found a highly significant positive correlation (p = 1.948^-20^; **Figure S1A**) between age and SDI. The same test was repeated on patient samples binned by age; <2 years or 2-4 years. SDI significantly increased with age over the first two years of life but not from 2-4 years of age (**Figure S1B**,**C; Table S1**). These associations are displayed using a linear model and 1-year age bins (**Figure 1A**). We used a linear model to test whether the basic demographics of sex and genotype might contribute to microbiome development, neither of which significantly impacted SDI (**Figure S2A**,**B**).

**Figure 1.**
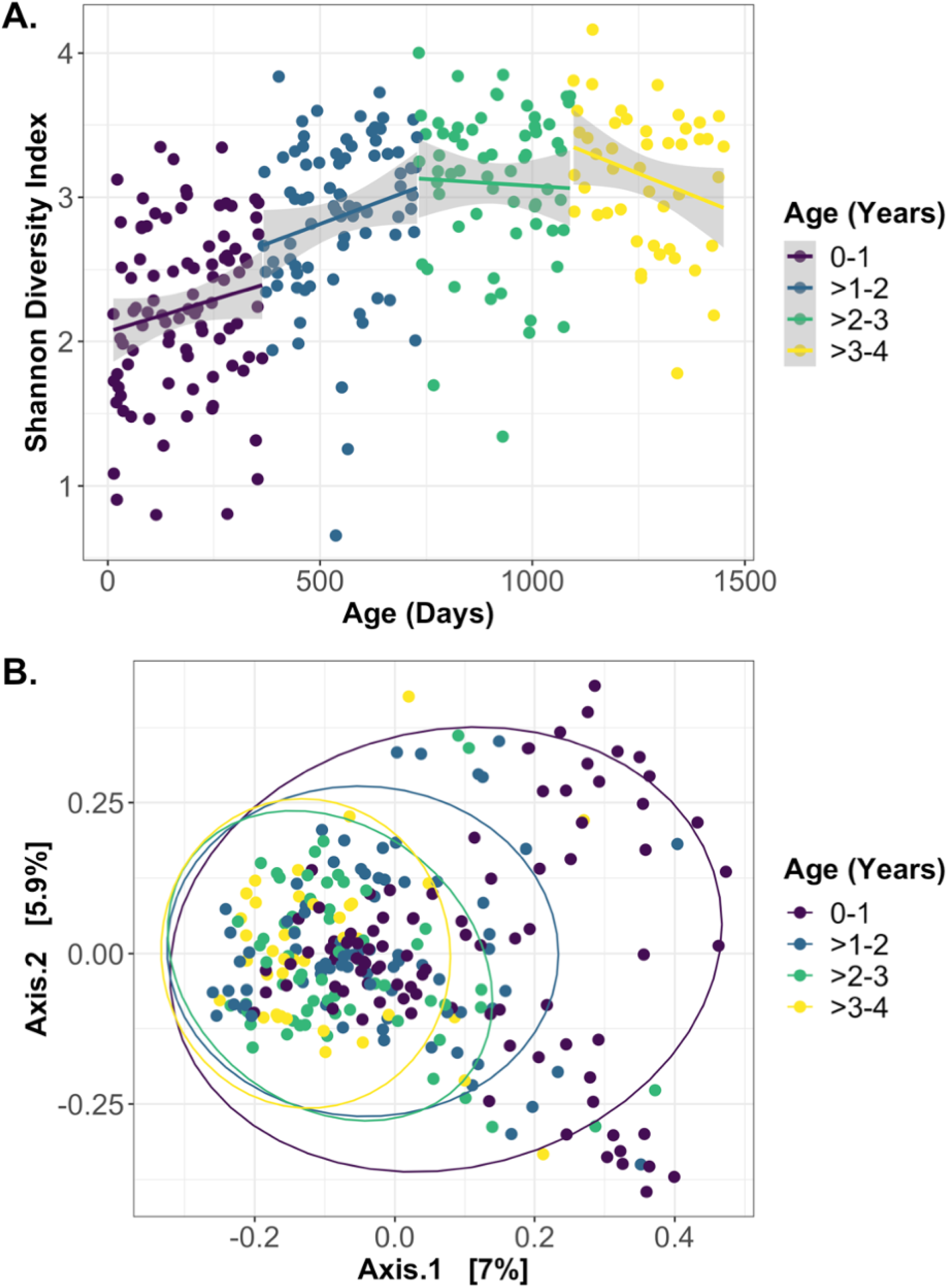
Alpha- and beta-diversity are age dependent. A) For each sample, patient age in days is graphed versus the Shannon Diversity Index (SDI). A linear model was used to visualize the increase in diversity over the first two years of life and the plateau of SDI above 2 years of age. B) Multidimensional Scaling (MDS) ordination of Bray-Curtis beta-diversity for each sample. Significance differences in beta-diversity were tested by PERMANOVA (permutational analysis of variance) for each pair of age bins, with all samples included in the model to adjust for additional time points. The strata function was used to adjust by individual patient for multiple sampling from the same cwCF. Samples from cwCF <1 year of age were significantly different (p<0.001) from all other time points. Samples from cwCF ages >1-2 were not significantly different from ages >2-3 (p=0.25), but were significantly different from >3-4 (p=0.01). Beta-diversity for samples from cwCF ages >2-3 and >3-4 were not significantly different (p = 0.18).

The Bray-Curtis distance metric was used to test the influence of age on overall microbiome composition, and significance was tested by PERMANOVA. The microbiome composition (beta-diversity) separation was strongest between samples <1 year of age relative to all other samples, which was significantly different from all other age-groups (**Figure 1B**). Interestingly, samples begin to cluster together more strongly after the first year of life. Consistent with SDI, sex and genotype did not significantly influence beta-diversity composition (**Figure S2C**,**D**). A shift in microbial composition is therefore apparent between the first and second year of life, with no significant impact of sex or genotype when controlling for patient.

### Large Alterations in the Microbiota Occur in Early Life

To understand the age-associated dynamics of the gut microbiome in cwCF, we first examined broad changes in relative abundance that occur over time at the phylum level. The ratio of Firmicutes/Bacteroidetes (F/B) has previously been associated with age and numerous health outcomes, including decreased F/B associated with inflammatory bowel disease (IBD) and conflicting F/B ratios linked to obesity [32-35]. In this cohort, the ratio of Firmicutes/Bacteroidetes increased significantly over the first year of life, but then plateaued from ages 1-4 years (**Figure 2A**). The Firmicutes/Bacteroidetes ratio was significantly negatively associated with Shannon Diversity Index overall, but only after the first year of life (**Figure 2B**).

**Figure 2.**
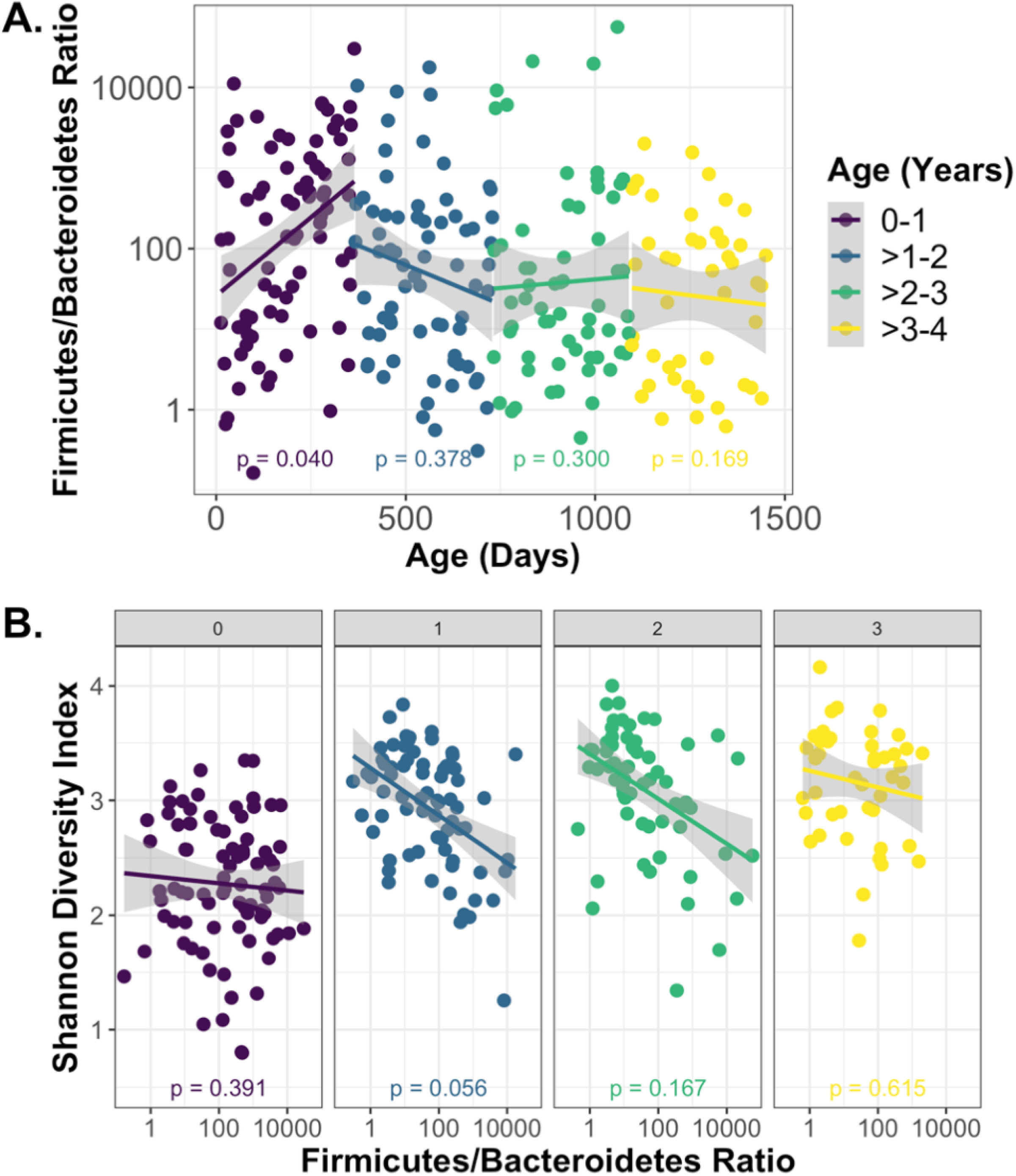
Firmicutes-to-Bacteroidetes ratios are age dependent. A) For each sample, patient age in days is graphed versus the Firmicutes/Bacteroidetes ratio. A linear model was used to visualize correlation between age and ratios for each year of life. Statistical significance was tested by mixed linear model. Firmicutes/Bacteroidetes was significantly correlated with age only in the first year of life but not after 1 year of age. B) Shannon Diversity Index is graphed versus the Firmicutes/Bacteroidetes ratio and the correlation is displayed by linear model. Each panel is representative of 1-year age bins (0: 0-1 years; 1: 1-2 years, 2: 2-3 years, 3: 3-4 years). The Firmicutes/Bacteroidetes ratio was significantly negatively associated with Shannon Diversity by mixed linear model for patients aged 1-4, but not for patients aged 0-1.

As cwCF age, major changes occur at the phylum level in early life and stabilize after approximately 500 days (**Figure 3A**). To examine these changes, we compared average phylum-level relative abundances of major taxa from all samples collected during the first 6 months (180 days) of life to all samples collected after 2 years (730 days) (**Table 1**; **Figure 3A**). These time points were chosen to best reflect the early microbiome compared to samples collected within the stable period from 2-4 years of age. Firmicutes undergo the largest overall change, increasing from an average relative abundance of 52% to an average relative abundance of 73% (**Figure 3A**). Proteobacteria and Verrucomicrobiota relative abundances also change unidirectionally. Proteobacteria begin at 24% average relative abundance during the first 6 months of life and reduce to 10%, while Verrucomicrobiota decrease from an initial level of 2% down to less than 0.1%. Bacteroidetes and Actinobacteria stabilize at 11% and 6%, at 4 years of life, respectively, but trend in opposite directions over time, with Bacteroidetes showing a modest increase after a reduction between 0 and ∼2 years of age.

**Table 1.** Relative abundance of major phyla for age-based subsets of samples.

| Phylum | All Samples | <6 months | >2 years |
| --- | --- | --- | --- |
| Firmicutes | 68.2 | 51.5 | 72.5 |
| Proteobacteria | 12.9 | 24.3 | 9.89 |
| Actinobacteria | 10.6 | 14.4 | 6.38 |
| Bacteroidetes | 7.85 | 7.51 | 11.1 |
| Verrucomicrobiota | 0.417 | 2.31 | 0.0660 |

**Figure 3.**
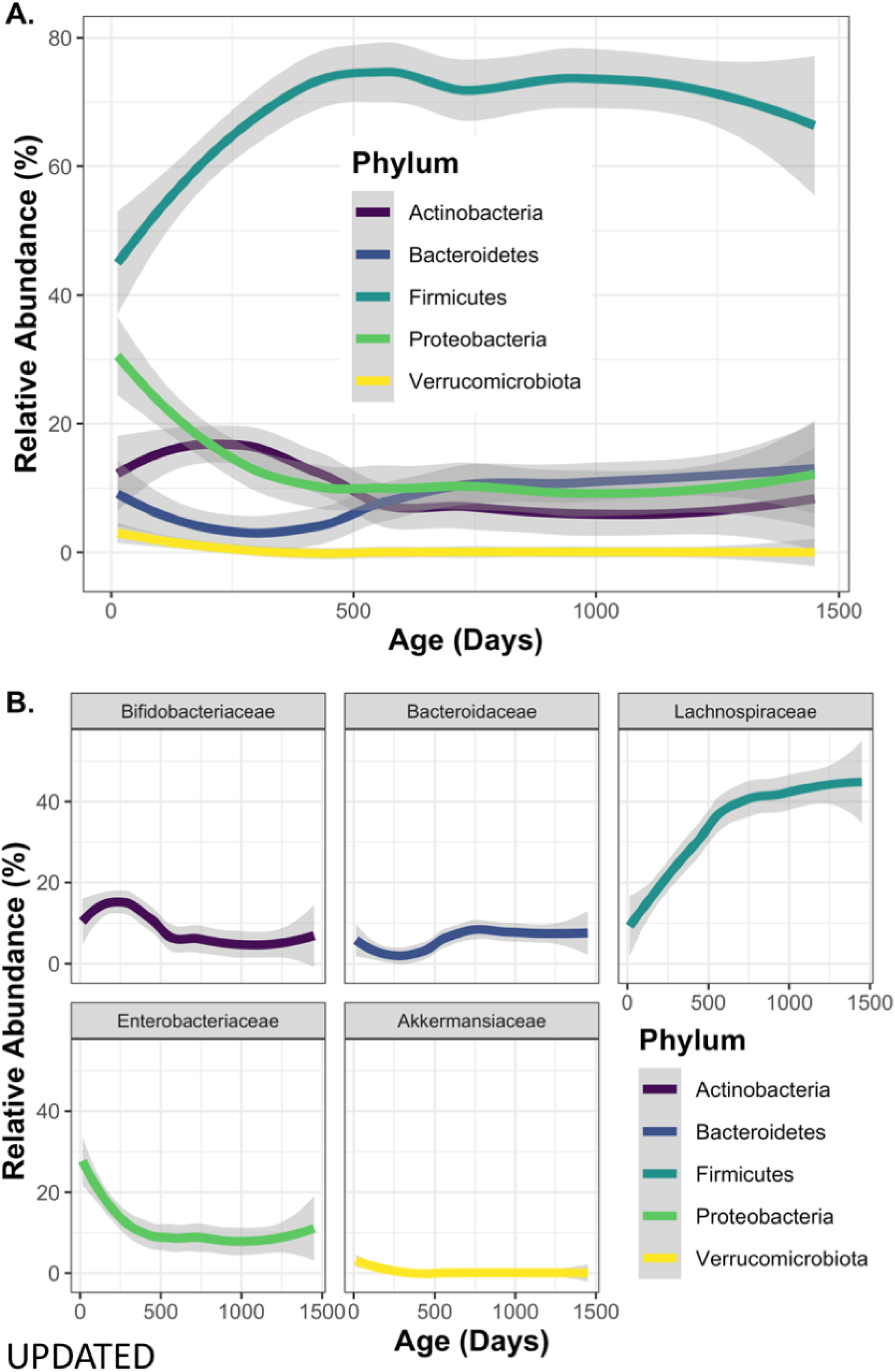
Microbial relative abundance changes with age. Age in days was graphed versus relative abundance of the indicated taxa for each sample and a linear model was used to visualize overall changes in microbial relative abundance at the A) phylum level and B) family level over the first four years of life.

The phyla Actinobacteria, Bacteroidetes, Proteobacteria, and Verrucomicrobiota are dominated by single families: Bifidobacteriaceae, Bacteroidaceae, Enterobacteriaceae, and Akkermansiaceae, respectively, and phylum level changes appear to be driven primarily by these taxa (**Figure 3B**; **Figure S3**). The largest individual family changes in Firmicutes are seen for Lachnospiraceae and changes in this family also reflects phylum-level changes (**Figure 3B**). However, Lachnospriraceae comprises <50% relative abundance of Firmicutes due to high diversity at the family level (**Figure S3**). These results highlight how the highest-abundance taxa change with age for cwCF. The overall patterns appear to be driven by a small number of bacterial families, especially within the Actinobacteria, Bacteroidetes, Proteobacteria, and Verrucomicrobiota.

### Changes in the Relative Abundance of Microbiota Over Time

We next determined the genera that changed significantly with age in our cohort. We compared a subset of samples grouped by age; samples <6 months and samples >2 years (**Figure 4A; Table S2**). Taxa that changed significantly with age were compared to relevant literature to determine whether changes moved more towards a ‘healthy-like’ composition or a ‘CF-like’ composition with age. This analysis has been applied such that taxa that are known to be higher in pwCF are labeled here as ‘more CF-like’ if the taxa increased with age and ‘less CF-like’ if decreased with age. For taxa known to be decreased in pwCF, those that increase with age are labeled as ‘less CF-like’ and ones that decrease with age become ‘more CF-like’. For example, *E. coli* is known to be increased in the gut microbiome of cwCF [8, 28]. We found that the *Escherichia/Shigella* group significantly decreased in relative abundance with increasing age, indicating that the relative abundance of *Escherichia/Shigella* shifts towards ‘healthy-like’ with age (**Figure S4A**). Notably, several taxa that are known to be decreased in pwCF, including butyrate-producers *Roseburia, Faecalibacterium, and Ruminococcus*, significantly increased with age in our cohort. The majority of taxa that changed significantly with age trended towards a ‘healthy-like’ composition over time. These results support the idea that the gut microbiome becomes less dysbiotic with age for pwCF, as has been observed in previous studies [8, 9]. However, the genus *Bacteroides*, which has reduced relative abundance in the gut microbiome of pwCF, does not significantly increase with age (**Figure S4B**) as has been reported in healthy cohorts. Additionally, four genera became more ‘CF-like’ with age, including: *Prevotella_7, Akkermansia, Bifidobacterium*, and *Blautia*. Of particular note is *Blautia*, which increases to >10% relative abundance in pwCF at >2 years of age (**Figure S4C**) and is the only taxa identified to both increase in relative abundance and become more CF-like with age. Interestingly, *Blautia* has both positive and negative associations with health in non-CF populations [36-39] but has rarely been discussed in the CF gut microbiome literature.

**Figure 4.**
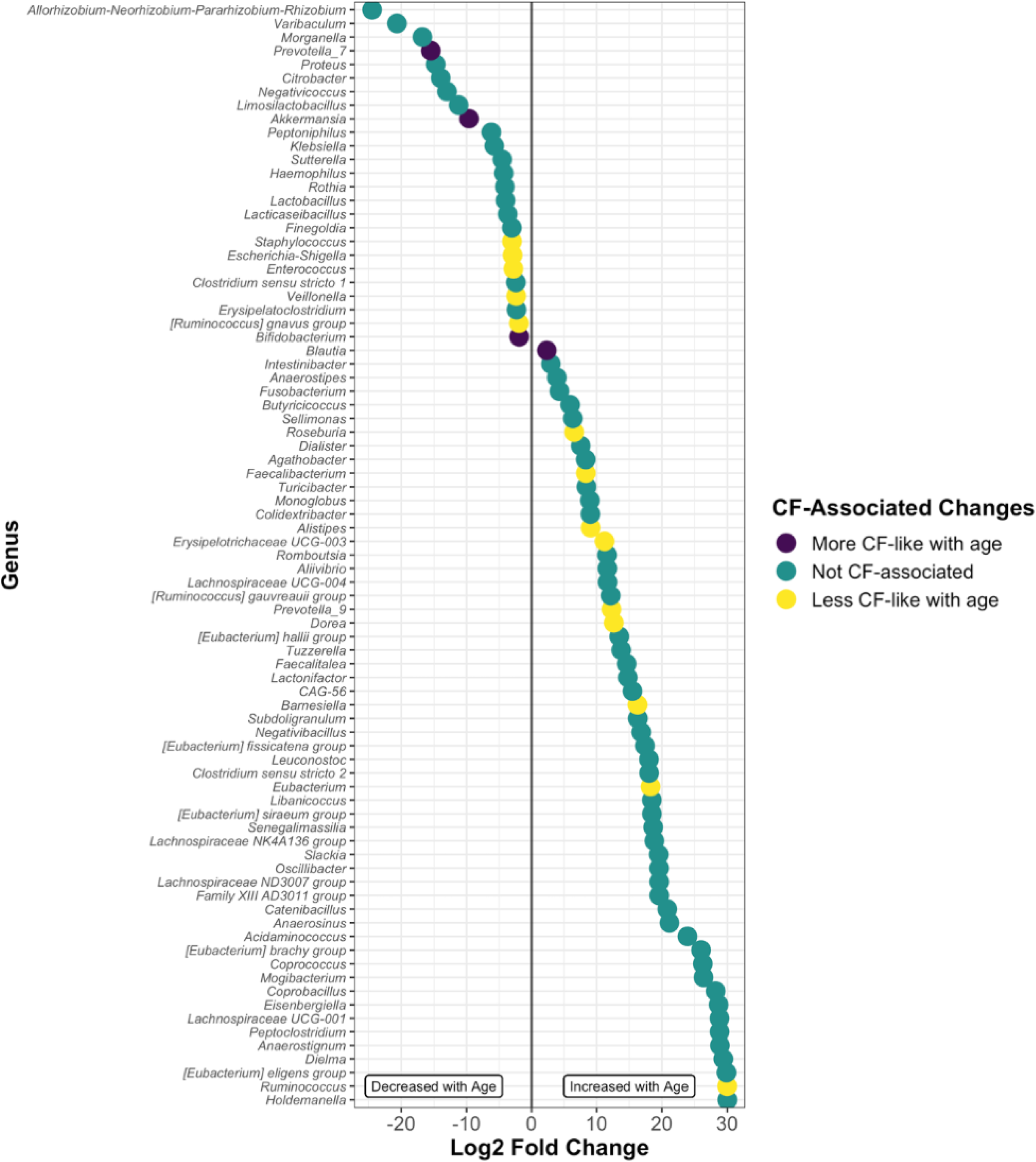
Significant alterations of abundance at the genus level occur with age. Log_2_ fold change of taxa that were significantly altered in samples from patients 2+ years of age relative to patients <6 months of age. Each dot represents a single genus and is color-coded by known CF-associated changes from current literature (see main text for details). Significance was determined by DESeq2.

### CF Lung Pathogens are Detected in the Gut

We examined each stool sample for both relative abundance and prevalence of opportunistic pathogens known to be important in CF, including *Gemella, Haemophilus, Neisseria, Prevotella, Pseudomonas, Staphylococcus, Stenotrophomonas, Streptococcus*, and *Veillonella* (**Figure 5A-B; Table S3**). Several of these genera are not known gut residents, so may be present transiently and indicate oropharyngeal seeding of the gut. However, of greater interest is the possibility that some of these taxa may be seeding the lungs from the gut. Previous work by Madan et al. demonstrated that several lung pathogens are detectable in stool prior to detection in the lungs [7].

**Figure 5.**
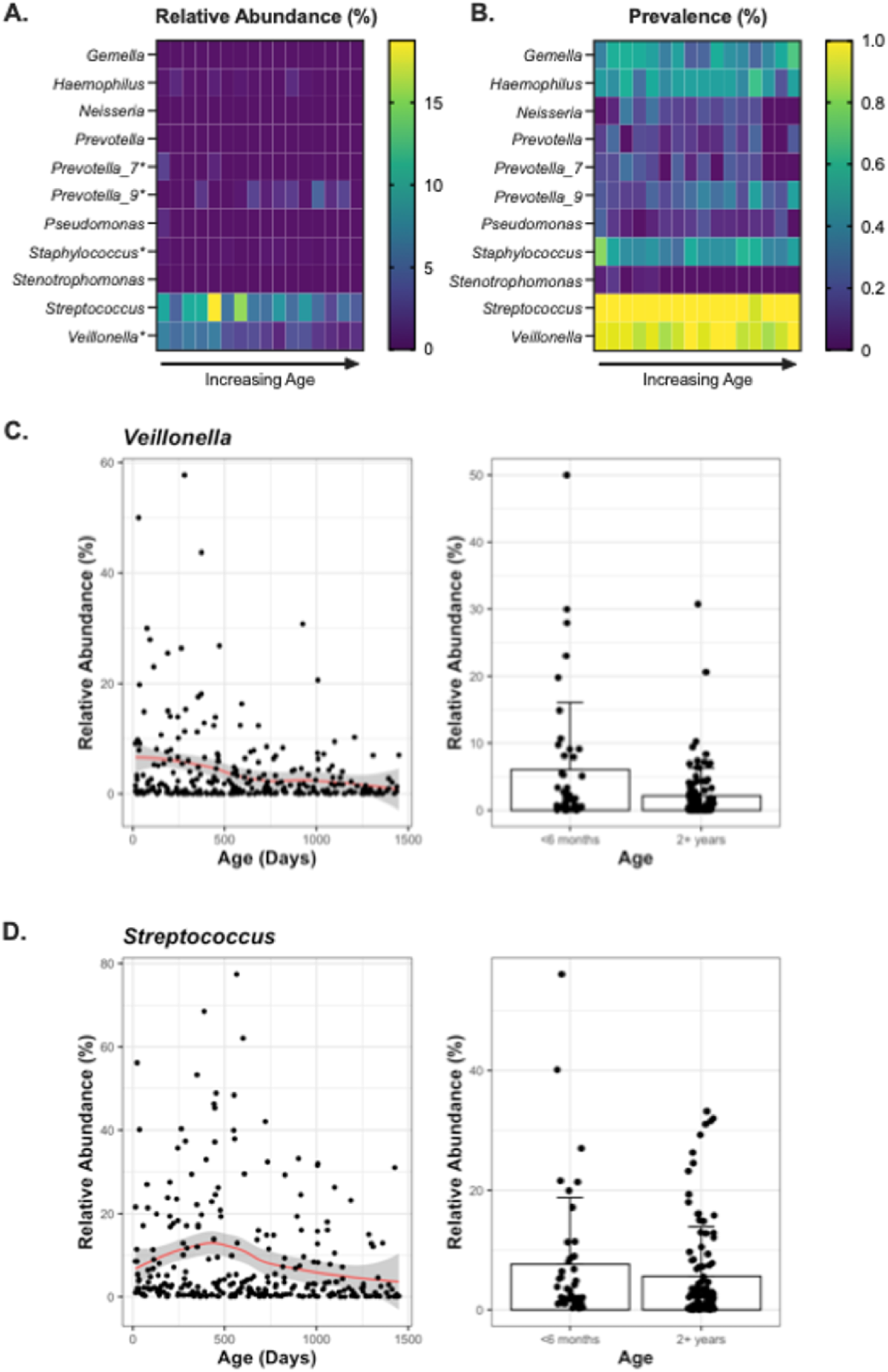
Age-related changes of genera associated with CF lung pathogenesis. A) Relative abundance and B) prevalence of each genus of interest is calculated for samples grouped by 3-month age bins. C-D) Changes over time of the relative abundance of *Veillonella* and *Streptococcus* are visualized by linear model (left) and the average relative abundance of is displayed for all samples from patients <6 months of age and 2+ years of age (right).

Three of the taxa examined (*Prevotella_7, Staphylococcus*, and *Veillonella*) were significantly higher in the first 6 months of life compared to >2 years of age, while *Prevotella_9* significantly increased in the older cwCF (**Figure 4-5**). *Streptococcus* and *Veillonella*, which are known to be gut resident microbes and have increased relative abundance in pwCF, had both the highest overall relative abundance and prevalence in the gut (**Figure 5; Table S3**). The relative abundance of *Veillonella*, but not *Streptococcus*, decreased significantly with age (**Figure 5C-D**). Next most prevalent (∼20-50%) were *Prevotella, Haemophilus, Gemella*, and *Staphylococcus*. Of these, *Staphylococcus* decreased a modestly but significantly with age. *Prevotella_7* and *Prevotella_9*, two unique ASVs within the Prevotellaceae family, also significantly changed with age, but *Prevotella_7* decreased with age while *Prevotella_9* increased with age. We observed lower rates of prevalence for *Stenotrophomonas, Pseudomonas*, and *Neisseria* at 1.8%, 9% and 14%, respectively and the relative abundances of these taxa do not change significantly with age. These data confirm that genera known to be important in CF lung disease are detected early and consistently in the gut microbiome of pwCF and could serve as a reservoir for seeding the lung.

### Early Crohn’s Dysbiosis Index Predicts Later Outcomes

Crohn’s disease is an intestinal inflammatory disorder with immunological and physiological symptoms overlapping with CF. The Crohn’s dysbiosis index (CDI) has previously been applied to gut 16S rRNA gene amplicon sequencing data from cohorts of pwCF as a measure of gut dysbiosis and was positively correlated with higher levels of the inflammatory marker fecal calprotectin [12]. We applied the CDI to our samples and found that few samples exceeded the “severe Crohn’s” metric of CD > 1 (**Figure 6A; Table S1**). Interestingly, when looking at the trends of cwCF individually, there did not appear to be a subset of patients with a consistently high CDI (**Figure S5A**). Instead, CDI correlated significantly with age, and most high CDI scores occurred during the first year of life, although not all cwCF had high scores at this age. CDI also significantly correlated with SDI; this is unsurprising given that microbiome diversity and age are highly intertwined (**Figure 6B**). However, the only other study to apply the CDI in pwCF did not find a correlation between SDI and CDI [12]; these differences are likely due to age differences between the cohorts.

**Figure 6.**
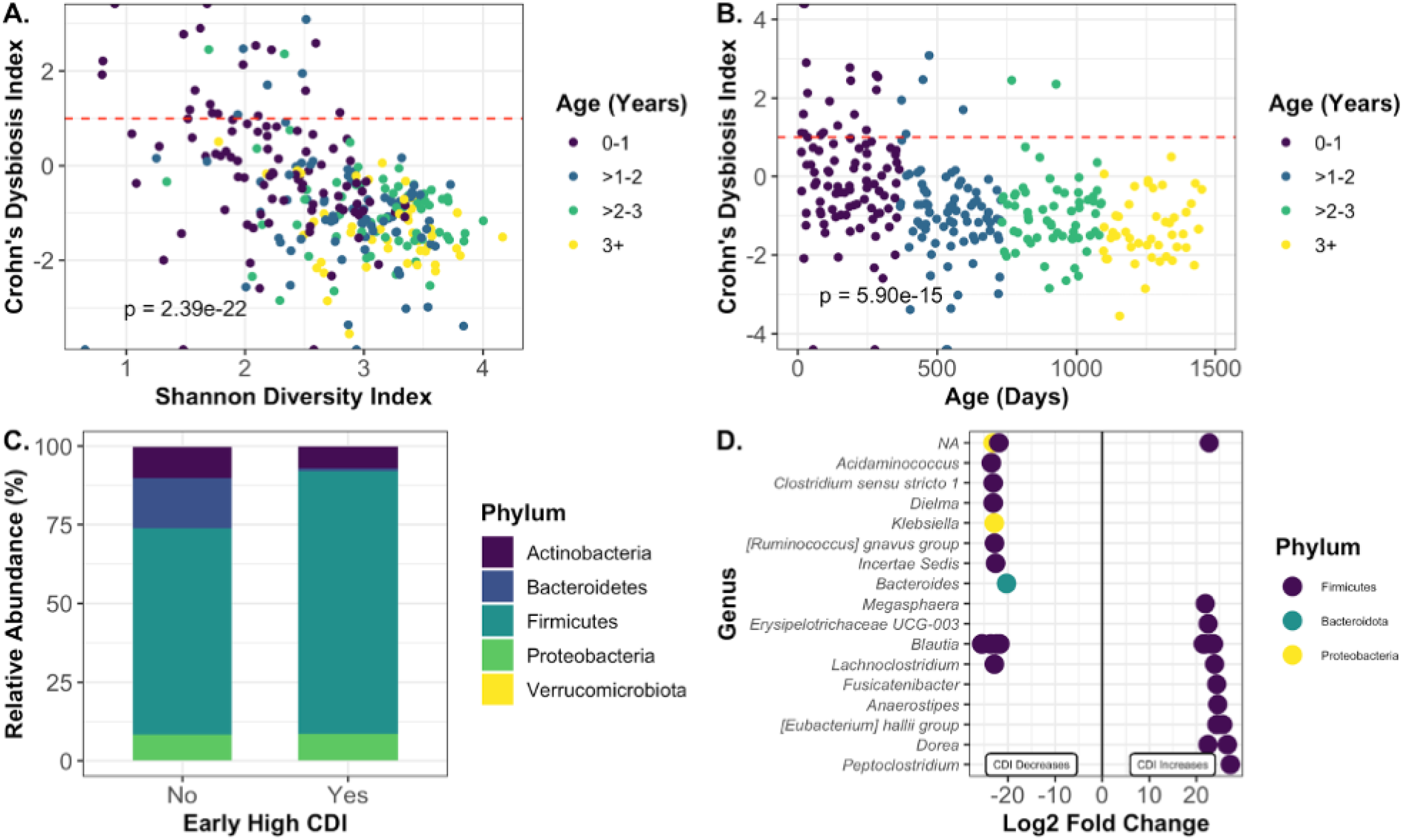
Early Crohn’s Dysbiosis Index is related to long-term changes in specific taxa. A) Shannon Diversity Index is plotted versus Crohn’s Dysbiosis Index for each sample. There was a significant inverse correlation between SDI and CDI as tested by mixed linear model. B) Age in days is plotted versus Crohn’s Dysbiosis Index for each sample. C-D) Samples from patients age 2 and above were grouped by whether the patient had an earlier sample with CDI > 0.5 (‘Yes’) or whether all earlier samples had a CDI < 0.5 (‘No’). C) Phylum-level relative abundances of samples with or without early high CDI. D) Log_2_ fold changes of ASVs that were significantly different in the early high dysbiosis group versus the early low dysbiosis group.

To determine whether a high CDI early in life could potentially shape later microbiome structure, we took samples from >2 years for all patients and classified them by whether or not the patient had at least one sample with a CDI > 0.5 during the first two years of life. Samples were excluded from the analysis if the patient did not have at least one sample before and after age 2. cwCF with samples from >2 years of age with a CDI >0.5 were also excluded, as they still have a high CDI score at this later time. A total of 20 cwCF had samples within these parameters, and a single sample from the latest collected time point was selected for each of these individuals. Samples were classified as having either ‘high’ (>0.5) or ‘low’ (<0.5) early CDI. Shannon diversity was not significantly different between the two groups, although it did trend higher in the early CDI group (**Figure S5B**). Interestingly, the gender distribution of the two groups was significantly different, with more males in the early high CDI group (**Figure S5C**). While this is a confounding factor that cannot be controlled for, it points to a potential influence of gender on early gut microbiome dysbiosis that is not reflected in overall gender-based differences in beta-diversity (**Figure S2**).

Analysis of taxa at the phylum level demonstrated that the relative abundance of Bacteroidetes was lower for patients who had a high early CDI (**Figure 6C; Table S4**). We used DESeq2 to determine significantly different ASVs between these groups (**Figure 6D; Table S4**). The overall decrease in Bacteroidetes at the phylum level is driven by a significant decrease in *Bacteroides*. However, not all of the changes were negative, as this group also saw a significant increase in the anti-inflammatory genus *Anaerostipes*. This finding highlights the impact that early microbiome disturbances may shape later outcomes, particularly for the genera *Bacteroides*.

## SUMMARY

We have performed a longitudinal analysis of microbiome samples from a cohort of 39 cwCF from birth through 4 years of age. For CF, which is a rare disease, this is a relatively large cohort with frequent longitudinal sampling and analysis of 281 total samples. We have demonstrated that the microbiome of cwCF follows some of the expected developmental patterns observed for infants and young children without CF, including increasing alpha diversity with age, although diversity plateaus ∼2 years of age for this CF cohort, and beta-diversity separation by age. An advantage of our cohort is the ability to leverage matched clinical data with the microbiome data. We have done that here with patient sex and genotype, neither of which significantly influence sample alpha-or beta-diversity in our cohort. These factors are known in the literature to influence gut microbial diversity and may be masked here by the influence of CF or by the cohort size.

Results from our analyses showed a marked difference between composition of the gut microbiome from 2-4 years of age when compared to infancy (< 6 months). This finding has several important clinical and translational implications given that the gut microbiota begins to stabilize and to resemble the adult profile beginning at about 2–3 years of age [31]. Such evidence signals the potential importance of modifying the microbiome of cwCF in early childhood. As diet changes significantly during this window of development, nutritional interventions could have far-reaching implications. Furthermore, this work raises the potential for a positive impact from CF-tailored probiotics.

Our findings have confirmed that many of the taxa known to be specifically increased or decreased in pwCF have the most extreme dysbiosis in early life and tend to move towards more typical abundances as cwCF age [8, 9]. A few microbes differ from this pattern, including *Akkermansia*, which is higher early in life in this cohort and is known to be decreased in pwCF. Furthermore, we note that *Bacteroides*, which is reduced in pwCF and may play an important role in immune programming, does not significantly increase with age. We have also confirmed the presence of many CF lung microbes in the gut of cwCF, highlighting the possibility of the lung being seeded directly from the gut early in life, consistent with previous findings.

Finally, we have applied the CDI to our cohort in a novel way. To date, the CDI has only been applied once in pwCF and never in a longitudinal dataset [12]. The CDI has previously been associated with calprotectin, a marker of intestinal inflammation, in both persons with Crohn’s disease and pwCF [12, 40]. These two diseases share many overlapping intestinal symptoms, and some tools from Crohn’s clinical diagnostics, such as the use of calprotectin as a marker of intestinal inflammation [41], have been used in recent CF studies [11, 20, 22, 28, 42, 43]. The CDI may therefore be another useful tool that can be employed to better understand the dynamics of the intestinal microbiome in pwCF and its relationship with clinical outcomes. Interestingly, we found that early high CDI appears to be associated with later microbiome structure (in years 3-4), particularly for *Bacteroides*. Previous work from our group and others demonstrated that *Bacteroides* is reduced in cwCF <1 year of age [4, 9, 11]. This current work extends that finding to later childhood and demonstrates that early perturbations can have long-term effects on *Bacteroides* relative abundance.

With this analysis, we are now able to better understand the development of the CF intestinal microbiome throughout early childhood. Future work within this cohort will focus on understanding how life history, including early exposure to antibiotics and breastmilk, impacts later microbiome structure, gastrointestinal and lung health outcomes. This analysis will help us to better understand how CF influences the microbiome and how microbiome development impacts the gut-lung axis.

## ACKNOWLEDGEMENTS

This work was supported by the Cystic Fibrosis Foundation (OTOOLE19GO) to G.A.O. and (MADAN18AO) to J.C.M and funded in part by US National Institute Health under award numbers T32-AI007363 to C.E.P and NIDCR R01-DE028154 to M.O.C. Additional support was provided by the Cystic Fibrosis Research Development Program (STANTO19R0) and DartCF (P30-DK117469).

## MATERIALS AND METHODS

### Patient population

We analyzed data from a cohort of 39 children with CF. Stool samples were collected longitudinally from 13 days up to 48 months of age (**Table 2**). A total of 281 stool samples were analyzed. Each patient had between 1-15 samples collected, with a median of 8 samples collected per patient. Age, sex, and genotype are also provided for each patient (**Table 2**).

**Table 2.** Patient and Sample Metadata.

| <u>Age</u> |  |  |  |  |
| --- | --- | --- | --- | --- |
| Age<br>(Years) | <1 | 1-2 | 2-3 | 3-4 |
| No. of<br>Samples | 93 | 80 | 62 | 46 |
| <u>Genotype</u> |  |  |  |  |
| Genotype: | dF508/dF508 | dF508/other | Rare/Rare | Not<br>Reported |
| No. of<br>Patients | 20 | 14 | 3 | 2 |
| No. of<br>Samples | 157 | 93 | 23 | 8 |
| <u>Sex</u> |  |  |  |  |
|  | Female | Male | Not<br>Reported |  |
| No. of<br>Patients | 20 | 17 | 2 |  |
| No. of<br>Samples | 158 | 115 | 8 |  |

### Sample collection and sequencing

Stool samples were collected by parents of cwCF and initially stored in a home freezer. Once samples arrived in the lab, they were stored at -80°C and later processed with Zymo fecal DNA miniprep kit (Cat #D6010). Paired-end reads were generated with 2×250 Illumina Miseq amplicon sequencing of the V4-V5 hypervariable region of the 16S rRNA gene (Woods Hole Marine Biological Laboratory). Raw data has been uploaded to the NCBI sequence read archive (SRA).

### 16S rDNA processing

Primer sequences were removed by CUTADAPT (v1.18). All subsequent pre-processing steps were performed in R version 3.6.0. Code is available at https://github.com/GeiselBiofilm. 39,577,426 raw paired-end reads were filtered and trimmed with dada2 version 1.14.1. Reads were then denoised, merged, and chimeras removed. The final counts were 30,765,091 total reads, 109,096 mean reads per sample, and 2,726 unique assigned sequence variants (ASVs). Taxonomy was assigned with DADA2 and the Silva v138.1 training set. See **Figure S6A** for a full analysis pipeline. ASV tables and sample information are available in the supplemental tables (**Table S1-2**).

### 16S rDNA analysis

All downstream analysis and visualization were performed in R (v4.0.2). Phyloseq (v1.32.0) and ggplot2 (v3.3.2) were used for data handling and visualization unless otherwise noted [44]. Reads per sample were graphed and filtered to include only samples with >10,000 reads (**Figure S6B**). One sample was removed because of low read count (<10^4^ reads) and the remaining 281 samples were further analyzed. DeSeq2 (v1.28.1) was used to call significantly differential abundances as described by McMurdie and Holmes [45, 46]. Linear mixed models were used for statistical regression analysis (nlme v3.1.151). For each sample, beta-diversity was calculated by Bray-Curtis distance, and multidimensional scaling (MDS) ordination was performed. Significant differences in beta-diversity were tested by permutational analysis of variance (PERMANOVA) (vegan v2.5.7).

### Crohn’s Dysbiosis Index

The CDI was applied to all samples as original described in Gevers, 2014 and first applied to CF in Enaud, 2019 [12, 40]. Briefly, counts of taxa associated with both positive and negative relative abundance changes in Crohn’s disease were summed for each sample. The log_10_ ratio of the [microbes increased in CD]/[microbes decreased in CD] was then calculated for each sample and associated with the relevant patient data.

### Accession Numbers

All sequence reads can be found in GenBank Sequence Read Archive with sequences found under accession number PRJNA170783 (SRP014429).

## Supplemental Figure Legends

**Figure S1. Shannon diversity index increases significantly with age**. A linear mixed effects model was used to determine that SDI increased significantly with age. ggPredict was used to visualize the linear regression model; both individual data points and the regression line are displayed above. A) Mixed linear regression of all samples. B) Mixed linear regression for samples in 2-year age bins from 0-2 years and >2-4 years. P-values are displayed in the graph.

**Figure S2. Alpha diversity and microbiome structure do not differ significantly based on gender or CFTR genotype**. A linear mixed effects model was used to determine that neither gender nor genotype led to significantly different Shannon Diversity Index (p >0.05). Data are visualized by linear model for A) gender and B) genotype. C-D) Bray-Curtis beta-diversity was calculated for each sample. Data were ordinated by multidimensional scaling (MDS) and plotted with color-coding by C) gender and D) genotype. Significance was tested by PERMANOVA and was not significantly different based on gender or genotype.

**Figure S3. Relative abundances of taxa at the family level**. Relative abundances of taxa at the family level are displayed for each phylum. Taxa with relative abundances within the phylum <10% are classified as other.

**Figure S4. Alterations of taxa with age**. For each panel, age versus genera-level relative abundances of samples is visualized by linear model (left) and the average relative abundances for samples from children with CF (cwCF) <6 months of age and cwCF 2+ years of age (right) are plotted for A) *Escherichia/Shigella* (p=5.13e-6), B) *Bacteroides* (p>0.05) and C) *Blautia* (p=3.33e-5). Significance was tested between samples from cwCF <6months of age versus 2+ years of age (right) by DESeq2 (Table S2).

**Figure S5. Crohn’s Dysbiosis Index**. A) Age versus Crohn’s Dysbiosis Index for each sample displayed by individual patient. The dashed red line is set at 1, the cutoff for severe Crohn’s in the original publication. B) Shannon Diversity Index and C) gender distribution for patients with (‘Yes’) or without (‘No’) an early high dysbiosis score. Differences in Shannon Diversity Index are not significant by student’s t-test (p>0.05), and differences in gender distribution are significant by chi-squared test (p<0.05).

**Figure S6. Microbiome analysis pipeline and data quality analysis**. A) Workflow of data processing steps for 16S rDNA gene amplicon sequencing analysis. B) Left: Amplicon sequence variants (ASVs) were ordered from most to least abundant, and total number of reads for each ASV is displayed. Right: Samples were ordered from those with the most counts to those with the fewest counts. A cutoff of 10,000 counts/sample was chosen, and as a consequence 1 sample was removed from the dataset.

## Supplemental Table Legends

**Table S1. Sample Data, Shannon Diversity Index, and Crohn’s Disease Index**.

**Table S2 Significantly Altered Taxa and All Assigned Sequence Variants**. A) Differential abundance was determined using DESeq2. B) Count table generated by DADA2 annotation pipeline.

**Table S3. Relative Abundances of CF Lung Pathogens Detected in Stool Samples**. A) Summary of Cystic Fibrosis (CF) pathogen relative abundance for all samples. B) CF pathogen relative abundance summarized by 3-month age groups.

**Table S4A. The Influence of Early CDI on Late Microbiota Composition**. A) Phylum Level average relative abundance for late samples from patients with and without early high Crohn’s Disease Index (CDI). B) ASVs detected as significantly different between late samples from patients with early low or early high CDI values.

